# MicroRNA-29a enhances and preserves stem-like CD8 T cell differentiation by regulating master epigenetic circuits of exhaustion

**DOI:** 10.64898/2026.05.25.727439

**Authors:** Xuebing Leng, Lance A. Buchness, Christine I. Rafie, Panagiotis I. Vlantis, Svetlana Ristin, Miguel A. Gallardo, Supipi L. Auwardt, Natasha K. Khatwani, Yuan-De Tan, Yuguang Ban, Corneliu M. Sologon, Benjamin B. Currall, Sion L. Williams, Kevin Van der Jeught, Jashodeep Datta, Alejandro Villarino, Aristeidis G. Telonis, Erietta Stelekati

## Abstract

CD8 T cells mediate protective immune responses. However, persisting antigens such as chronic viruses or tumors redirect CD8 T cell differentiation to a sub-optimal, epigenetically defined state called exhaustion. Exhausted T cells (T_EX_) lose their ability to persist long-term and to initiate functional memory responses. Checkpoint inhibitor blockade temporarily restores effector functions, but immune reinvigoration is not long-lasting, due to the epigenetic stability of T_EX_. Therefore, epigenetic reprogramming of T_EX_ leading to durable T cell responses is essential to improve disease control. Here, we demonstrate that a single microRNA (miR), miR-29a, epigenetically re-directs T_EX_ differentiation and preserves T_EX_ into a stem-like state, leading to long-term persisting progenitor T_EX_. MiR-29a rewires epigenetic maintenance programs, including downregulation of key exhaustion-associated regulators (Dnmt1, Dnmt3a, and Dnmt3b), alongside increased expression of progenitor- and stemness-associated genes such as Tcf7 and Il7r. These reprogrammed CD8 T cells are more sensitive to PD-L1 checkpoint blockade. Ectopic expression of miR-29a combined with aPD-L1 treatments enhances effector responses, while preserving T cell stemness. Together, our findings suggest that miR-29a can be leveraged to overcome current barriers to immune checkpoint blockade.

**Highlights:**

- MiR-29a rewires key exhaustion-associated epigenetic maintenance programs, while enhancing stemness-associated transcriptional circuits.
- MiR-29a drives extensive remodeling of accessible chromatin in T_EX_.
- MiR-29a preserves newly generated progenitor T_EX_ in a durable, epigenetically defined stem-like state with increased effector function.
- MiR-29a synergizes with aPD-L1; while miR-29a preserves progenitor T_EX_ state, addition of aPD-L1 enhances the cytotoxic potential of these progenitor T_EX_ cells.

## Introduction

CD8 T cells orchestrate anti-viral and anti-tumor immune responses (Zhang and Bevan, 2011). However, chronic infections and cancer cause CD8 T cell exhaustion. T cell exhaustion is a sub-optimal differentiation state with unique epigenetic identity, characterized by the inability of CD8 T cells to provide immunological protection (Blank et al., 2019; McLane et al., 2019; Pauken and Wherry, 2015; Scott-Browne et al., 2016; Wherry and Kurachi, 2015). Improving the function of exhausted T cells (T_EX_) by antagonizing checkpoint inhibitors, such as PD-1, is a potent immunotherapeutic strategy (Sharma et al., 2021, 2023). However, a considerable portion of patients develop resistance to immunotherapy that is often attributed to the epigenetic stability of T_EX_ as a state that cannot be reversed (Abdel-Hakeem et al., 2021; Belk et al., 2022; Pauken et al., 2016; Restifo et al., 2016; Sen et al., 2016; Tonnerre et al., 2021; Yates et al., 2021; Zebley et al., 2020). Therefore, to overcome the development of resistance, novel strategies need to be developed to epigenetically reprogram T_EX_ and achieve a durable response.

Transcriptionally, T_EX_ are characterized by the regulation of diverse biological processes (Bengsch et al., 2018; Giles et al., 2022; Yao et al., 2019; Zebley et al., 2020). Thus, targeting a single pathway is unlikely to provide full T_EX_ reinvigoration by rewiring the epigenetic regulatory networks that provide robustness to the state of exhaustion (Philip et al., 2017; Scott-Browne et al., 2016; Sen et al., 2016; Yates et al., 2021; Zebley et al., 2020). In this context, microRNAs (miRs) are promising candidates with a profound impact on the epigenetic and transcriptional regulation by simultaneously targeting several RNAs within multiple pathways (Gagnon and Ansel, 2019; Garzon et al., 2010; O’Connell et al., 2010; Rupaimoole and Slack, 2017; Wells et al., 2020). We recently identified miR-29a as a unique memory-associated miR that is repressed in T_EX_ (Stelekati et al., 2022). Using the prototypical mouse model of exhaustion induced by chronic viral Lymphocytic Choriomeningitis Virus (LCMV) infection, we demonstrated that ectopic expression of miR-29a significantly enhances T_EX_ persistence, attenuates exhaustion, and promotes differentiation of a T_EX_ subset with progenitor, stem-like characteristics. Here, we demonstrate that miR-29a is a key regulatory hub in resolving the state of exhaustion and that it can enhance the effects of immunotherapy to promote CD8 T cell responses. We demonstrate that miR-29a transcriptionally and epigenetically reprograms T_EX_ differentiation resulting in durable, persisting progenitor T_EX_ with preserved stem-like properties and increased effector function.

## Results

We induced exhaustion in C57BL/6 mice by infecting them with a chronic viral strain of LCMV (clone 13; cl-13) (Barber et al., 2006; Stelekati et al., 2022). Ectopic overexpression (OE) of miR-29a was induced in TCR-transgenic CD8 T cells (P14) recognizing the immunodominant epitope of LCMV (gp^33-41^). Retrovirally transduced miR-29a OE or control transduced P14 cells were adoptively transferred to congenically marked recipient mice infected with LCMV cl-13 (**Figure 1A**). Host mice were subsequently treated with aPD-L1 or control isotype and transferred P14 cells were analyzed in the spleens upon cessation of treatments (**Figure 1B**). MiR-29a enhanced the numbers of adoptively transferred CD8 T cells (**Figure 1C**), consistent with our previous findings (Stelekati et al., 2022). This effect was not due to alterations in the viral loads or an overall increase in total CD8 T cells, as viral loads and host CD8 T cells were not affected (**Figure S1A-B**). The combination of miR-29a OE and aPD-L1 further increased the numbers of P14 cells (**Figure 1C**), suggesting a synergistic effect between miR-29a and aPD-L1 in retaining the T_EX_ numbers in the context of persistent antigen.

**Figure 1.**
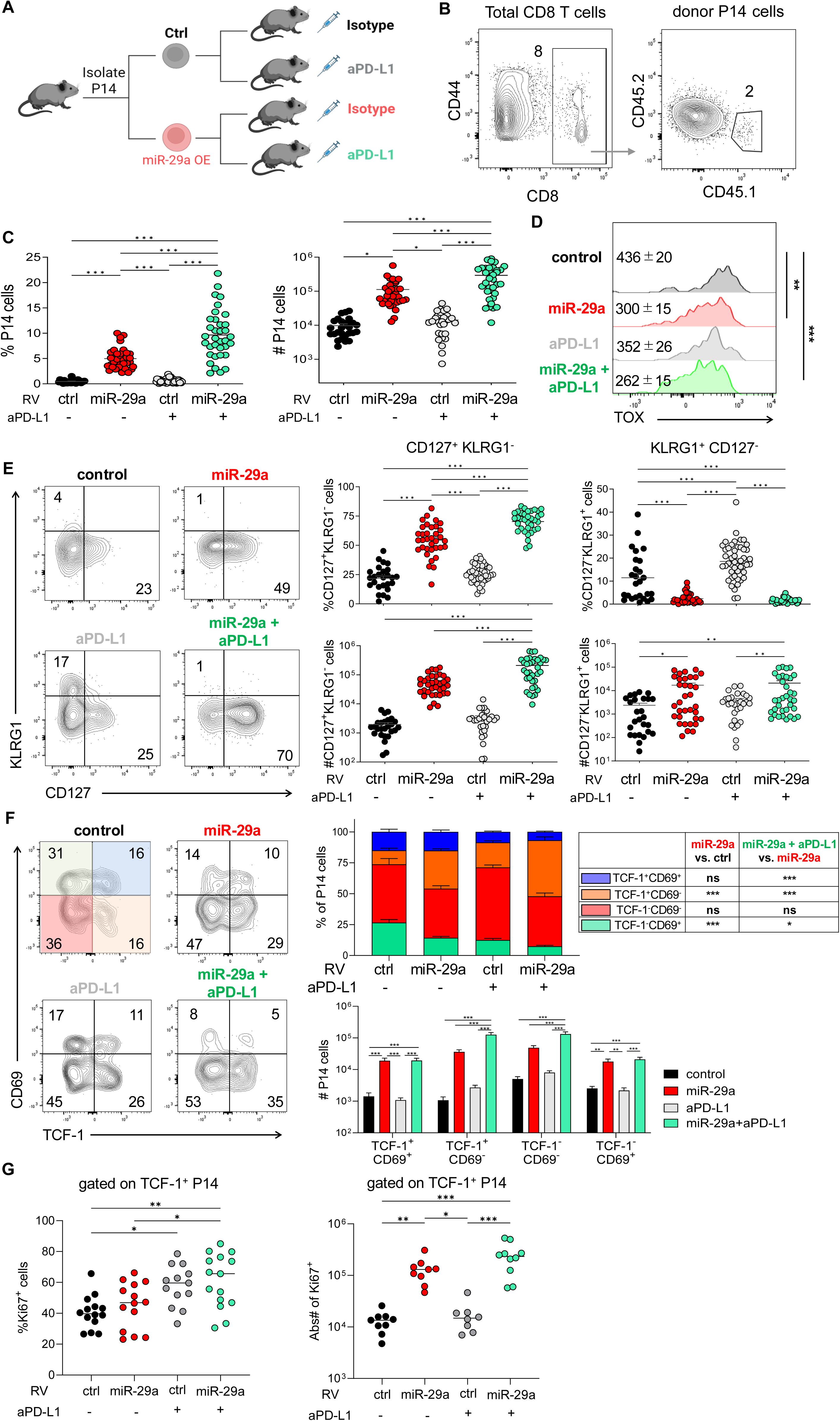
Combination of miR-29a and aPD-L1 promotes progenitor T_EX_ differentiation with increased proliferative potential. **(A)** P14 cells were transduced with miR-29a OE or control retrovirus, sorted for VEX^+^ expression and transferred into recipient congenically marked mice that were infected with LCMV cl-13 at 48 hrs earlier. Isotype or anti-PD-L1 (200 µg/injection) was administered every third day starting at d21 p.i., for a total of 5 doses. **(B)** Gating strategy for donor P14 cells. **(C)** Quantification of donor P14 cells in the spleens at d35 p.i. **(D)** TOX expression in donor P14 cells at d35 p.i. Numbers represent mean±SEM of mean fluorescence intensity (MFI) of TOX in P14 cells. **(E)** CD127^+^KLRG1^-^ and CD127^-^KLRG1^+^ P14 cells were quantified in spleens at d35 p.i. **(F)** T_EX_ P14 subsets were identified by TCF-1 and CD69 expression. **(G)** Ki-67 expression was evaluated in TCF-1^+^ P14 cells. Data are pooled from 5 independent experiments with n>8. FACS plots are gated on donor VEX^+^ P14 cells and numbers represent percentages out of total donor VEX^+^ P14 cells. Results are indicated as mean ± SEM. Statistical analysis was performed using One-way ANOVA with Tukey’s multiple comparisons (\**P* < 0.05, \*\**P* < 0.01, and \*\*\**P* < 0.001). Ctrl, control RV. Illustration in **(A)** created using BioRender.com.

Therefore, we investigated the potential synergism between miR-29a and aPD-L1 on T_EX_ differentiation. T_EX_ differentiation is directed by the key transcription factor TOX (Alfei et al., 2019; Khan et al., 2019; Scott et al., 2019; Seo et al., 2019; Yao et al., 2019). We found that miR-29a OE reduced TOX expression (**Figure 1D**) and expression of inhibitory molecules (**Figure S1C**) while retaining surface expression of PD-1, suggesting that miR-29a OE cells retain high levels of activation in response to TCR stimulation but attenuate signaling pathways leading to exhaustion. Addition of aPD-L1 did not further reduce TOX or inhibitory receptor expression compared to miR-29a OE, suggesting that miR-29a regulates T_EX_ differentiation more potently than aPD-L1.

Persistent TCR stimulation during chronic infection is detrimental for the differentiation of memory T cells (T_MEM_) from T_MEM_-precursor CD8 T cells (CD127^+^KLRG1^-^ cells) (Chang et al., 2014; Pipkin and Rao, 2009; Staron et al., 2014; Tonnerre et al., 2021). Reinvigoration by aPD-L1 primarily promotes the differentiation of short-lived effector-like CD8 T cells, thus, immune reinvigoration is not long-term sustained. We analyzed the dynamics of these cell populations and found that miR-29a promoted the differentiation of T_MEM_-precursor CD8 T cells consistent with a role for miR-29a in promoting T_MEM_ (Yee Mon et al., 2021). Combination with aPD-L1 further enhanced the proportion and numbers of these precursors; instead aPD-L1 only enhanced effector-like (KLRG1^+^CD127^-^) cells (**Figure 1E**). Importantly, miR-29a did not inhibit the differentiation of effector-like CD8 T cells, as the absolute numbers of miR-29a OE KLRG1^+^CD127^-^ CD8 T cells were also increased compared to control, presumably due to greater accumulation of total P14 cells.

T_EX_ is a heterogeneous pool of cells (Beltra et al., 2020; Hudson et al., 2019; Zander et al., 2019). A progenitor, stem-like T_EX_ subset, that retains proliferative potential and responsiveness to immunotherapy is governed by the transcription factor TCF-1 (Chen et al., 2019b; Hashimoto et al., 2022; He et al., 2016b; Im et al., 2016; Leong et al., 2016; Utzschneider et al., 2016; Zhou et al., 2010). MiR-29a increased progenitor TCF-1^+^ subset differentiation and combination of miR-29a OE and aPD-L1 further promoted the TCF-1^+^ subset that lacks expression of CD69, a circulating, highly proliferative T_EX_ subset (**Figure 1F**). In contrast, combination of miR-29a OE and aPD-L1 proportionally reduced the TCF-1^-^ subset that expresses CD69, characterized as the most terminally differentiated, resident T_EX_ subset (**Figure 1F**). To understand whether the altered T_EX_ subset differentiation was due to altered proliferation, we examined Ki-67 expression within T_EX_ subsets. As expected, aPD-L1 increased the proliferation of TCF-1^+^ cells (**Figure 1G**) but not TCF-1^-^ cells (**Figure S1D**). MiR-29a enhanced the proliferation of TCF-1^+^ cells only in the presence of aPD-L1 and preserved a significantly higher number of proliferating TCF-1^+^ cells (**Figure 1G**). These results suggest that while aPD-L1 increases the proliferation of progenitor T_EX_, miR-29a enhances their durability and persistence, therefore preserves the pool of proliferating T_EX_.

With the robust effects of miR-29a in T_EX_ differentiation and dynamics at hand, we wanted to understand the transcriptional changes induced by miR-29a in combination with aPD-L1. To this end, we analyzed the transcriptome of donor cells at day 35 p.i. (from **Figure 1A**). MiR-29a had a robust effect on the transcriptome, while addition of aPD-L1 only moderately altered the transcriptional profile of T_EX_ (**Figure 2A**). The transcriptional profile of miR-29a OE cells correlated with our previous findings (**Figure S2A**) and anti-correlated with the geneset of the miR-29a predicted mRNA targets (**Figure S2B**), suggesting that the miR-29a effects are, at least partially, due to direct mRNA targeting. More importantly, although aPD-L1 alone had no significant impact on the expression of miR-29a predicted mRNA targets, the addition of aPD-L1 to miR-29a OE de-enriched for miR-29a predicted targets signature, suggesting that the aPD-L1 effects were not driven by direct mRNA targeting (**Figure S2B**). MiR-29a induced downregulation of expected transcripts as compared to control (**Figure 2B**), consistent with the role of miR-29a promoting quiescence and a stem-like T_EX_ state. Interestingly, addition of aPD-L1 only moderately increased the number of downregulated transcripts and decreased the number of upregulated transcripts (**Figure 2C**), suggesting that the synergistic effect of miR-29a and aPD-L1 in promoting stem-like T_EX_ differentiation may reflect a less transcriptionally active state.

**Figure 2.**
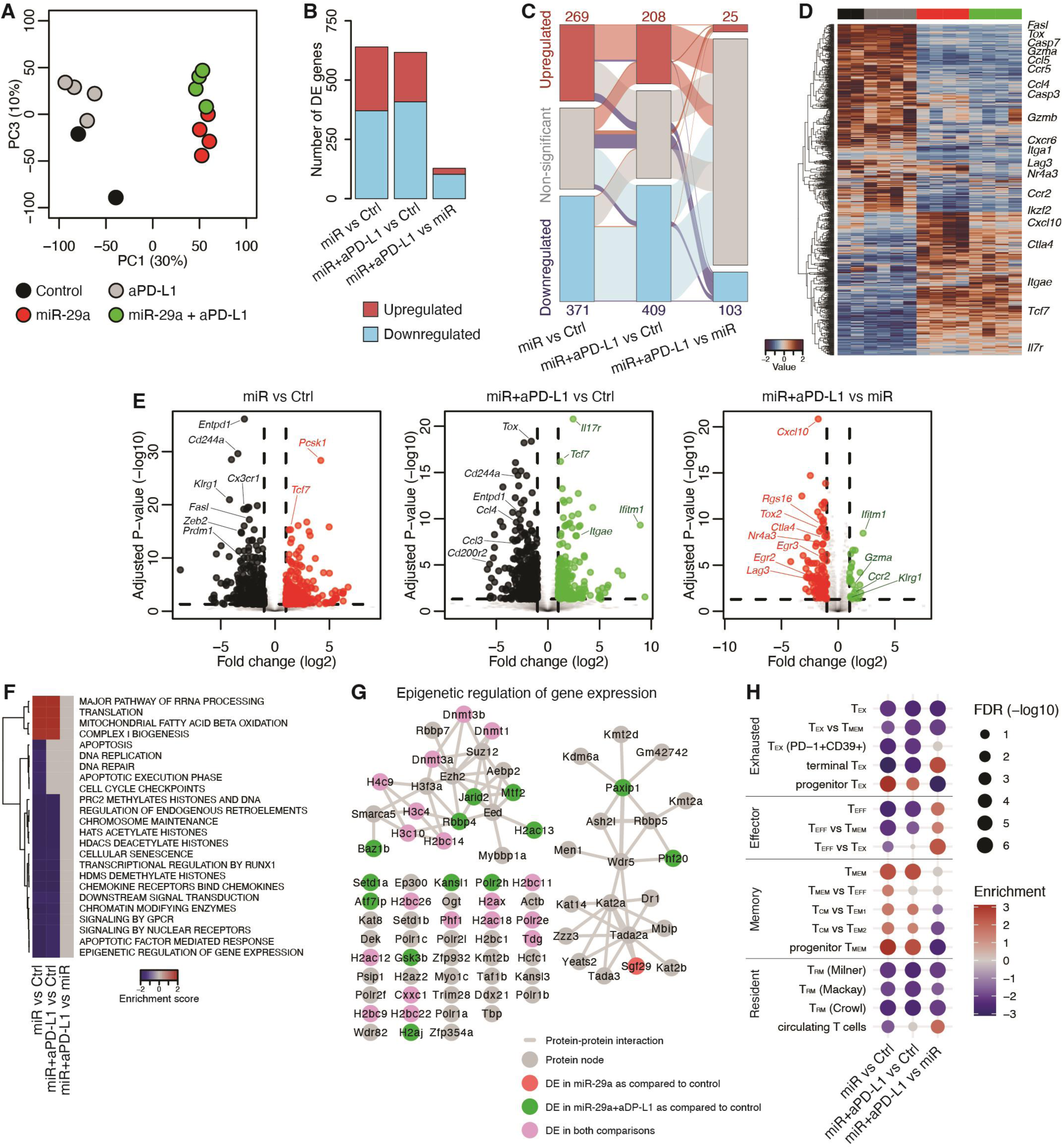
MiR-29a promotes memory-like transcriptional signatures, while aPD-L1 enhances effector-like transcriptional programs. P14 cells were transduced with miR-29a OE or control retrovirus, sorted for VEX^+^ expression and transferred into congenically marked recipient mice that were infected with LCMV cl-13 at 48 hrs earlier. Isotype or aPD-L1 (200 µg/injection) was administered every third day starting at d21 p.i., for a total of 5 doses. VEX^+^ P14 cells were sorted from spleens at d35 p.i. and RNA Sequencing was performed. **(A)** Principal Component Analysis (PCA) **(B-D)** Barplot (b) alluvial plot (c) and heatmap (d) showing the number, overlaps and normalized expression values of the differentially expressed genes (FDR<5%; absolute log_2_ fold change >1 per DESeq2) among the indicated comparisons. **(E)** Volcano plots of the indicated comparisons. **(F)** heatmap showing the significantly enriched Reactome pathways in the differentially expressed genes (FDR<10% per GSEA). **(G)** Protein-protein interaction network depicting the interactions among the proteins of the epigenetic regulation of gene expression pathway. Nodes are colored by whether the expression of the gene was found to be significantly different after miR-29 OE with or without aPD-L1 treatment. **(H)** Bubble plot showing the enrichment of T cell expression signatures in the differentially expressed genes after miR-29a OE with or without aPD-L1 treatment. The size of the value is inversely proportional to the FDR and the color to the enrichment score as per GSEA.

MiR-29a upregulated genes known to attenuate exhaustion and promote stem-like T_EX_ differentiation, including *Tcf7* and *Il7r* (**Figure 2D**), confirming our immunophenotyping (**Figure 1**). In contrast, key transcription factors known to regulate effector and T_EX_ differentiation (*Tox, Prdm1, Zeb2*) (Dominguez et al., 2015; Kallies et al., 2009; Omilusik et al., 2015; Rutishauser et al., 2009; Shin et al., 2009) were downregulated by miR-29a (**Figure 2D-E**), consistent with a role of miR-29a in promoting stem-like differentiation and diverting away from terminal differentiation. However, addition of aPD-L1 upregulated some effector-related transcripts (*Klrg1, Gzma*), further attenuated inhibitory receptors (*Ctla4, Lag3*) and inhibited key antigen-driven transcription factors downstream of TCR activation (*Nr4a3, Tox2, Egr2, Egr3*) (Chen et al., 2019a; Jung et al., 2022; Miao et al., 2017) (**Figure 2E**). These results suggest that, while miR-29a promotes progenitor T_EX_ differentiation, addition of aPD-L1 plays a supportive role in repressing expression of transcription factors dependent on TCR stimulation. In line with these results, miR-29a downregulated a transcriptional signature for response to antigen (**Figure S2C**). In contrast, aPD-L1 enriched for the antigen response signature compared to miR-29a OE alone, as antagonizing the PD-1 pathway releases the immunological break of TCR signaling. At the pathway level, miR-29a caused the downregulation of apoptosis-related pathways, arguing for a role for miR-29a in enhancing T_EX_ persistence and durability (**Figure 2F**). Interestingly, multiple epigenetic-related pathways were downregulated by miR-29a suggesting a role for miR-29a in regulating the epigenetics of exhaustion. In fact, a global pathway of epigenetic regulation of gene expression was downregulated in miR-29a OE P14 cells with key epigenetic regulators of exhaustion (*Dnmt1, Dnmt3a, Dnmt3b*) driving this enrichment (**Figure 2G**). Among the few pathways induced by miR-29a were those implicated in mitochondrial metabolism and protein translation (**Figure 2F**). Addition of aPD-L1 did not significantly affect the pathways implicated in cellular differentiation, apart from inducing de-enrichment for apoptosis- and cell cycle-related pathways, suggesting that, while miR-29a promotes a less transcriptionally active yet persisting cell state, combination of miR-29a and aPD-L1 may promote a more effector-like state.

Therefore, we asked whether combination of miR-29a and aPD-L1 altered the known differentiation trajectory of T_EX._ We found that miR-29a OE cells were depleted from T_EX_-related transcriptional signatures, confirming our immunophenotyping, and addition of aPD-L1 did not significantly alter this negative enrichment for exhaustion-related signatures (**Figure 2H**). Furthermore, miR-29a OE P14 cells enriched for stem- and progenitor-like transcriptional signatures and depleted for effector-like and terminally differentiated gene signatures. Combination of aPD-L1 with miR-29a increased enrichment for effector-like and terminal signatures and depletion for stem-like and memory-like signatures (**Figure 2H**). MiR-29a also depleted cells from resident memory (T_RM_) transcriptional signatures, while addition of aPD-L1 enriched for a circulating transcriptional signature (**Figure 2H**). Together, these results suggest that miR-29a preserves a durable, stem-like, functional cell state despite chronic stimulation, while combination of miR-29a with aPD-L1 enhances effector-like T cell differentiation and diverts away from stem-like T cell states, consistent with the role of aPD-L1 in promoting proliferation and effector-like differentiation.

Since miR-29a transcriptionally regulated epigenetic pathways and the known T_EX_ subset differentiation trajectory, we asked whether miR-29a epigenetically altered T cell differentiation. Thus, we analyzed the open chromatin profile of donor cells at day 35 p.i. Strikingly, miR-29a had a robust effect on epigenetic regulation of T_EX_ differentiation (**Figure 3A**), although addition of aPD-L1 only moderately affected the epigenetics of T_EX_, as expected, due to the known inefficiency of aPD-L1 to induce complete T_EX_ epigenetic rewiring (Pauken et al., 2016). MiR-29a induced an increase in accessibility in 5,533 regions (**Figure 3B-D**) These regions overlapped with pseudogenes (**Figure 3E**) and were enriched in intergenic regions (**Figure 3F**). These results suggest the opening of regulatory regions, including distal enhancers and looping regions, rather than of specific molecular processes. Interestingly, miR-29a reduced accessibility of 4,030 regions (**Figure 3C**) that are strongly enriched for intronic elements (**Figure 3F**), indicating a repression of intragenic enhancers and reduced transcriptional activity, coinciding with a reduced transcriptional activation (**Figure 2B**). Intersecting these regions with previously defined epigenetic alterations of T cell exhaustion revealed that miR-29a has a significant reinvigorating effect and is associated with epigenetic recovery of T_EX_ (**Figure S2D-E).** We then performed transcription factor motif analysis and found motifs for key transcription factors known to modulate T cell differentiation enriched in chromatin regions gained by miR-29a. Notably, the binding motifs for *Lef1* and *Tcf7*, two transcription factors regulating T cell stemness, were both enriched in regions gained by miR-29a (**Figure 3G**), consistent with a role for miR-29a in promoting the progenitor T_EX_ subset. Key transcription factor binding motifs, including interacting AP-1/NFAT factors, were enriched in regions gained by miR-29a but lost by combination of miR-29a and aPD-L1, highlighting the distinct effects of miR-29a and aPD-L1; while miR-29a promotes stem-like, progenitor states, aPD-L1 promotes effector-like differentiation. Surprisingly, we observed an enrichment in transcription factor motifs in regions gained by miR-29a, such as Fosl2, BATF, Runx1 and IRF, for which we also observed transcriptional downregulation (**Table S1**). Thus, we suggest miR-29a post-transcriptionally regulates key transcription factors downstream of TCR activation to instruct T cell differentiation. In line with a role for miR-29a in regulating T cell differentiation at the epigenetic level, we found that open chromatin peaks that were lost upon miR-29a OE, were enriched for effector- and terminal exhaustion-related genesets, while addition of aPD-L1 did not further alter the T_EX_ differentiation trajectory epigenetically (**Figure 3H**).

**Figure 3.**
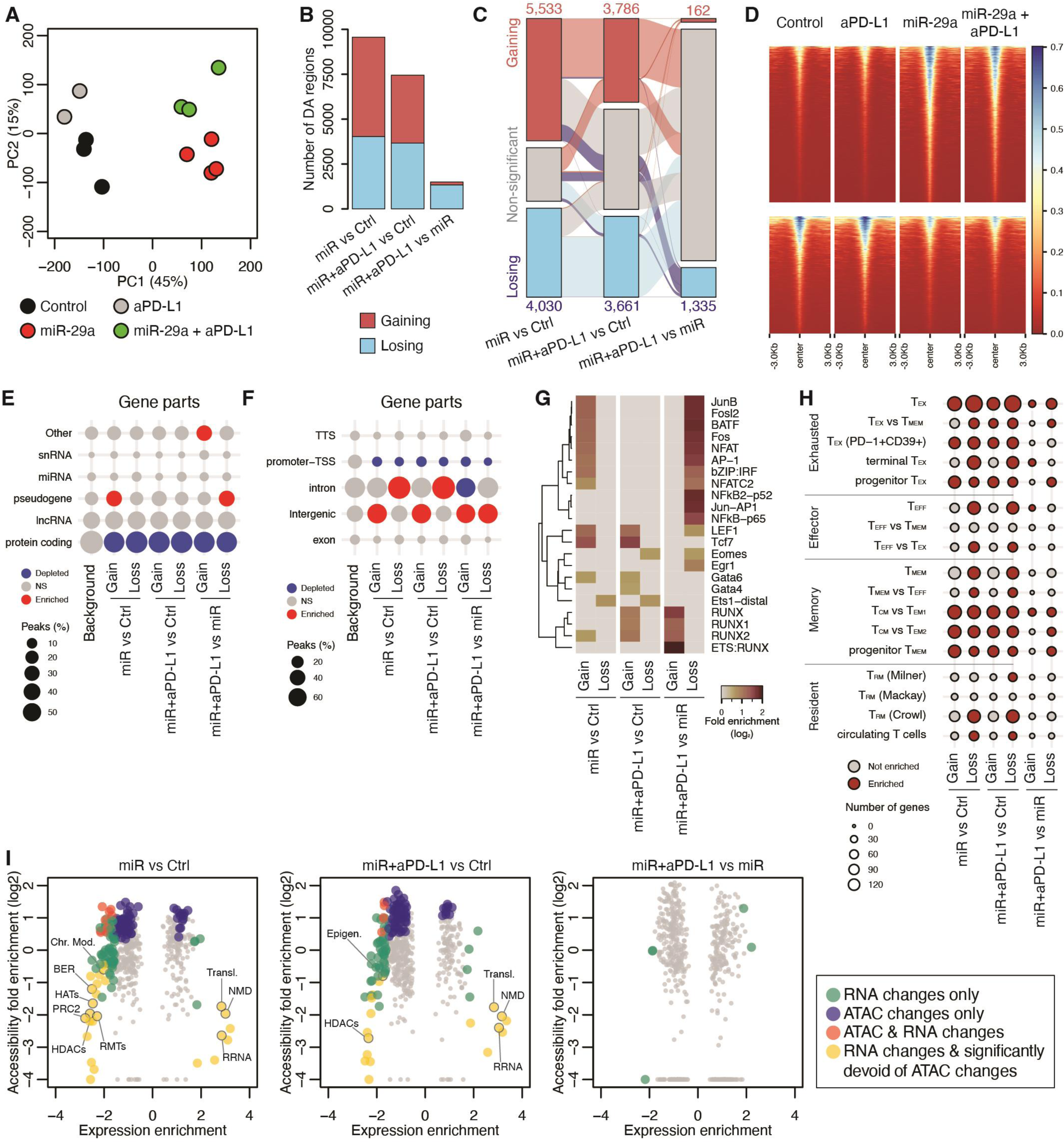
MiR-29a epigenetically alters T_EX_ differentiation. **(A)** Principal Component Analysis (PCA) of the DNA accessibility of P14 cells transduced with miR-29a OE or control retrovirus, transferred to LCMV-infected mice and treated with isotype or aPD-L1. **(B-D)**, Barplot (b) alluvial plot (c) and heatmap (d) showing the number, overlaps and accessibility, respectively of the differential ATAC peaks (FDR<5%; absolute log_2_ fold change > 1 as per DESeq2) among the indicated comparisons. **(E-F)**, Bubble plots showing the distribution of the background and differential ATAC peaks in gene types (e) or gene parts (f). The size is proportional to the percentage of peaks overlapping each category while the color indicates significant enrichment or depletion (P-value<0.05 and >5% difference compared to the background, as per chi-squared tests). **(G)** Heatmap showing the enrichment of transcription factor binding motifs in the peaks with significant accessibility changes. Shown are only selected motifs; the full list of motifs is shown in **Table S1**. **(H)** Bubble plot showing the enrichment of T cell expression signatures in the genes overlapping significant accessibility changes. The size of the bubble is proportional to the number of genes overlapping each gene set. Red color indicates significant enrichment (FDR<10% as per hypergeometric tests). **(I)** Scatter plots showing the enrichment of biological processes in expression (X axis) or accessibility (Y axis) change. Each dot represents a Reactome pathway, the X axis notes the enrichment score as per GSEA, the Y axis shows the fold enrichment of the pathway in the set of genes overlapping accessibility change (union of peaks significantly gaining or losing accessibility signal). Pathways are colored by significant RNA (FDR<10% as per GSEA) or accessibility changes (FDR<10%, log_2_ fold enrichment > 1 as per hypergeometric test). Chr. Mod: chromatin modifying enzymes; BER: base excision repair; HATs: histone acetyltransferases; HDACs: histone deacetylaces; PRC2: polycmob repressive complex; RMTs: histone argine methyltransferase; Transl.: translation; NMD: nonsense-mediated decay; RRNA: ribosomal RNA processing; Epigen.: epigenetic regulation of gene expression.

With both RNA and ATAC data at hand, we examined which of the differentially expressed biological processes are also enriched in accessibility changes. We reasoned that any discordance between expression and accessibility would suggest post-transcriptional regulation, possibly by miR-29a. Focusing on the pathways that are significantly enriched in expression changes but significantly devoid of accessibility changes at the respective genes, we found that miR-29 OE leads to expression-specific downregulation of a series of epigenetic pathways, including histone acetylases, histone deacetylases and histone arginine methyltransferases. We noted that the miR-29a OE also led to the expression-specific upregulation processes associated with overall expression output, including translation, ribosomal RNA processing and non-sense mediated decay (**Figure 3I**). These data indicate that miR-29a targets core gene sets regulating cell identity maintenance and primes the transcriptome towards biosynthetic processes and proliferation.

T_EX_ reinvigoration provided by aPD-L1 therapy is not longitudinal, creating the need for extended and recurrent cycles of therapy, potentially enhancing toxicities. Since miR-29a OE enhanced progenitor T_EX_ durability and persistence, we hypothesized that the miR-29a effects could persist long-term, in contrast to the aPD-L1 effects (**Figure S3A**). As expected, the effects of aPD-L1 were undetectable at day 100, i.e. two months post-treatment (**Figure S3B)**. In contrast, the effects of miR-29a OE were still detectable two months post-treatment, with significantly increased numbers of P14 cells, reinstating the role of miR-29a in redirecting T_EX_ differentiation to long-term persisting cell states. The CD8 T cells persisting at day 100 retained a T_MEM_-precursor phenotype with low inhibitory receptor and TOX expression and contained higher numbers of progenitor (TCF-1^+^ CD69^-^) T_EX_ subsets (**Figure S3C-F**). Together, this data suggests that miR-29a drives long-term persisting stem-like T_EX_ differentiation with T_MEM_-precursor phenotypes.

The robust and sustained effects of miR-29a on T_EX_ differentiation and the regulation of progenitor T_EX_ persistence led us to hypothesize that miR-29a may fundamentally alter progenitor T_EX_ differentiation instead of only increasing their numbers. Thus, we differentiated miR-29a OE or control T_EX_ in LCMV cl-13 infected hosts (**Figure 4A**) and at day 21 p.i. we isolated progenitor T_EX_ (Ly108^+^CX3CR1^-^) and terminal T_EX_ (Ly108^-^PD1^+^) (**Figure 4B**). We then transferred equal numbers of progenitor or terminal T_EX_ either miR-29a-OE or control to infection-matched secondary recipients, which were then treated with aPD-L1 or isotype control. Despite transfer of equal numbers, progenitor miR-29a OE P14 cells exhibited a more robust expansion responding to aPD-L1 therapy compared to controls (**Figure 4C**). Interestingly, however, miR-29a OE did not enhance persistence of terminal T_EX_ cells (**Figure 4C**), suggesting that miR-29a primarily affects progenitor T_EX_. The increased numbers of progenitor miR-29a-OE T_EX_ was not due to increased proliferation (**Figure 4D**), which led us to hypothesize that miR-29a intrinsically altered the long-term persistence and durability of progenitor T_EX_. Indeed, despite similar progenitor-like phenotype at transfer, after 2 weeks upon secondary adoptive transfer, miR-29a OE progenitor T_EX_ retained a T_EX_ progenitor phenotype, while control cells acquired a more terminal effector-like phenotype (**Figure 4E**). Importantly, while miR-29a preserved a progenitor-like T_EX_ state, it also increased the capacity of these progenitor T_EX_ cells to produce cytotoxic molecules, including IFN-γ and Granzyme B (**Figure 4F**). Collectively, these results suggest that miR-29a intrinsically alters progenitor T_EX_ differentiation, thus, enhances persistence of T_EX_ and responsiveness to aPD-L1.

**Figure 4.**
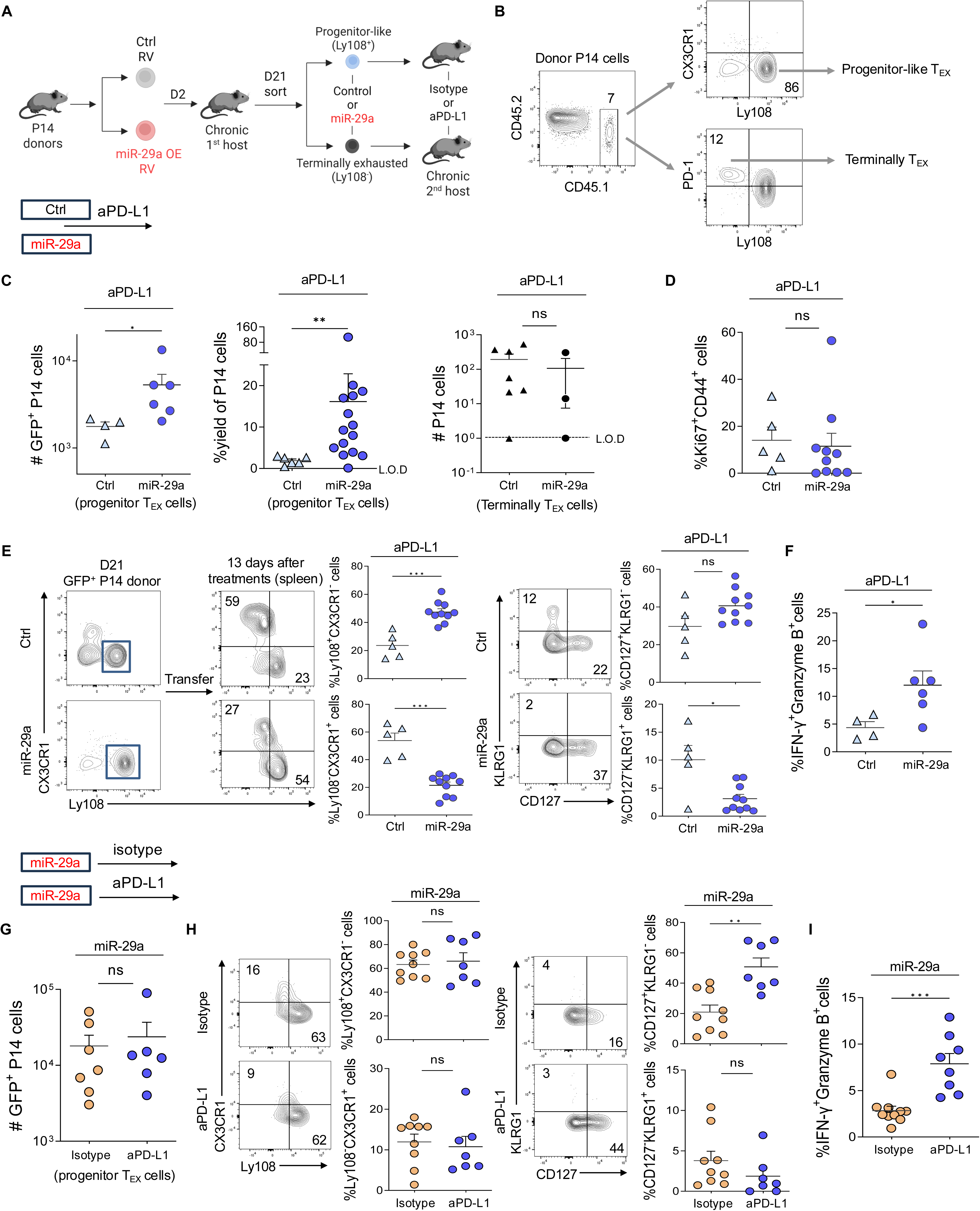
MiR-29a preserves stemness in progenitor-like T_EX_ while increasing their effector functions upon anti-PD-L1. **(A-B)** P14 cells were transduced with miR-29a OE or control retrovirus, sorted for GFP^+^ expression and transferred into congenically marked recipient mice that were infected with LCMV cl-13 at 48 hrs earlier. At day 21 p.i., progenitor-like T_EX_ (Ly108^+^ CX3CR1^-^) P14 cells and terminal T_EX_ (Ly108^-^ PD-1^+^) P14 cells were sorted and equal numbers (9x10^4^ cells) were transferred to infection-matched congenically marked recipients. Secondary recipients were treated with isotype or aPD-L1, as in Figure 1A. Transferred P14 cells were analyzed in spleens of secondary recipients at 2 wks after transfer. **(C)** Numbers of miR-29a OE GFP^+^ P14 cells and ctrl GFP^+^ P14 cells in secondary host mice upon aPD-L1 treatment. **(D)** Ki-67 expression in GFP^+^ P14 cells**. (E)** Representative FACS plots showing gating strategy at d21 and the phenotype of transferred P14 cells at 2 wks after transfer. Frequency (left) of Ly108^+^CX3CR1^-^ and Ly108^-^CX3CR1^+^ GFP^+^ P14 cells from **(B).**Frequency (right) of CD127^+^KLRG1^-^ and CD127^-^KLRG1^+^ GFP^+^ P14 cells from **(B).. (F)** Frequency of IFN-γ^+^Granzyme B^+^ GFP^+^ P14 cells from **(B). (G)** Numbers of miR-29a OE GFP^+^ P14 cells in secondary host mice upon aPD-L1 or isotype treatment. **(H)** Phenotype of GFP^+^ P14 cells from **(G). (I)** Frequency of IFN-γ^+^Granzyme B^+^ GFP^+^ P14 cells from **(G).** Data are pooled from 2 independent experiments with n>5 **(B-I)**. FACS plots are gated on donor GFP^+^ P14 cells and numbers represent percentages out of total donor GFP^+^ P14 cells. Results are indicated as mean ± SEM. Statistical analysis is performed using Mann-Whitney test (\**P* < 0.05, \*\**P* < 0.01, and \*\*\**P* < 0.001). Ctrl, control RV. Illustration in **(A)** created using BioRender.com.

We then asked whether the effects of miR-29a were independent of aPD-L1 treatment or if aPD-L1 treatment reinforced the miR-29a effects. The effects of miR-29a in enhancing progenitor T_EX_ persistence and durability and retaining a progenitor T_EX_ phenotype were largely independent of aPD-L1 treatments (**Figure 4G-H**). However, combination of aPD-L1 with miR-29a OE enhanced the capacity of progenitor T_EX_ cells to produce IFN-γ (**Figure 4I; Figure S4**). These results suggest the synergistic effects of combining miR-29a with aPD-L1 treatments; while miR-29a increases the numbers of proliferating T_EX_ and preserves a long-term persisting progenitor T_EX_ state, addition of aPD-L1 enhances the cytotoxic potential of these progenitor T_EX_ cells.

## Discussion

Our data highlight that miR-29a epigenetically reprograms T_EX_ differentiation and combined with aPD-L1 can modulate progenitor-like T cells. Strikingly, miR-29a sustains the proliferative capacity and progenitor-like phenotype of T_EX_ even after aPD-L1 treatment withdrawal, while restoring their effector functions, suggesting that miR-29a predominantly drives these durable effects. The combination of miR-29a and aPD-L1 not only enhances the durability of progenitor-like T cells but also preserves them in a stem-like state while increasing their cytokine production. Our study suggests a novel strategy to enhance the durability of progenitor-like T_EX_ and regulate T_EX_ differentiation to improve current immunotherapies.

The stable epigenetic profile of T_EX_ is a major obstacle for inducing durable T cell reinvigoration upon immunotherapy. Here, we demonstrate robust epigenetic changes induced during T_EX_ differentiation by regulating miR-29a expression. MiR-29a increased accessibility in intergenic gene regions not innately accessible in canonical T_EX_, suggesting increased accessibility for various enhancers, silencers, insulators, and other distal genomic regulatory elements (He et al., 2016a; Sun and Dong, 2025). MiR-29a epigenetically preserved stemness in T_EX_ by increasing chromatin accessibility for known stem-like associated transcription factors, including *Tcf7* and *Lef1* (Chen et al., 2019b, 2026). Interestingly, processed-pseudogene regions, known to play an important role in regulating stemness, self-renewal, and pluripotency (Poliseno et al., 2024), also became more accessible with miR-29a. In contrast, miR-29a antagonized the T_EX_ epigenetic landscape, as demonstrated by the loss of chromatin access for inhibitory receptors. The capacity of miR-29a to promote stem-like fate at the epigenetic level suggests durable maintenance of the T_EX_ subsets responding to checkpoint blockade.

Previous studies have defined the heterogeneity of T_EX_, with the progenitor-like T_EX_ subset playing an indispensable role in maintaining the pool of T_EX_ (Beltra et al., 2020; Hudson et al., 2019; Utzschneider et al., 2020; Zander et al., 2019) and mediating responses to checkpoint inhibitor therapy (Hashimoto et al., 2022; Im et al., 2016). Promoting the differentiation of progenitor-like T_EX_ is a current promising strategy to regulate T_EX_ differentiation and improve responses to immunotherapies. For example, IL-2 cytokine therapy has been shown to drive TCF-1^+^ stem-like T_EX_ toward an effector CD8 T cell phenotype in synergy with anti-PD-1 therapy (Hashimoto et al., 2022). Our study demonstrates the potential to not only enhance the proliferation and promote the differentiation of canonical progenitor-like T_EX_, but to further preserve newly generated progenitor T_EX_ in a durable, epigenetically defined stem-like state with increased effector function, by regulating the expression levels of miR-29a. These findings will guide the development of novel immunotherapeutics (such as nanoparticle-based *in vivo* delivery of miR-29a, in combination with anti-PD-1 therapy) and the design of next-generation CAR T cells with increased durability and retained effector function.

## Acknowledgements

Research reported herein was performed in part at the Onco-Genomics Shared Resource (RRID: SCR_022502) and at the Flow Cytometry Shared Resource (RRID: SCR022501) at Sylvester Comprehensive Cancer Center, which is supported by the National Cancer Institute (NCI; P30CA240139). The content is solely the responsibility of the authors and does not necessarily represent the official views of the NIH. We thank Dr. Oliver Umland at the DRI flow cytometry core facility of the University of Miami Miller School of Medicine for helping with flow cytometric analysis and cell sorting. We thank Nathaniel Knudsen for helping with python programming for gene mapper pro v.1. This study was supported by NIH/NIAID 1R01AI183292-01(E.S.), NIH/NIAID R21AI178184 (E.S.), ACS-IRG (E.S.), B+ Foundation (E.S.). C.I.R. is supported by NIH 5F31CA294908-02.

## Author contributions

X.L., L.A.B., A.G.T., and E.S. designed the overall study. X.L., L.A.B., C.I.R., S.R., M.A.G., S.L.A., and N.K.K. performed the experiments. A.G.T. and P.I.V. led the transcriptomics and epigenetic studies. C.M.S., B.B.C and S.L.W. performed RNA and ATAC Sequencing. Y.T., Y.B. and A.V. contributed with transcriptomic data analysis. K.V. der J., J.D., and A.V. participated in study design and interpretation of data. X.L., L.A.B., A.G.T., and E.S. wrote the manuscript. All co-authors commented on, participated in writing and approved the manuscript.

## Methods

### Key Resources Table

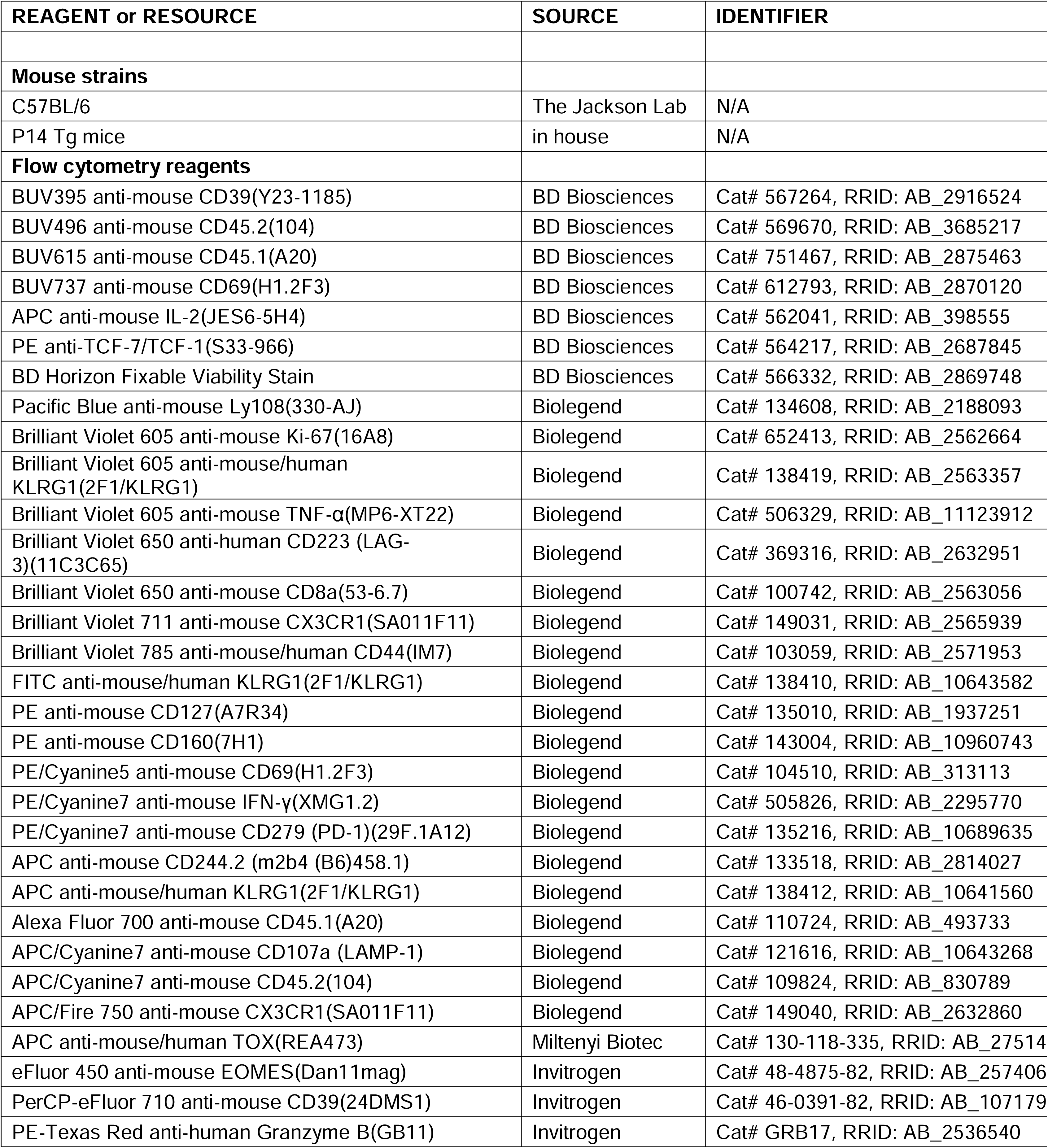
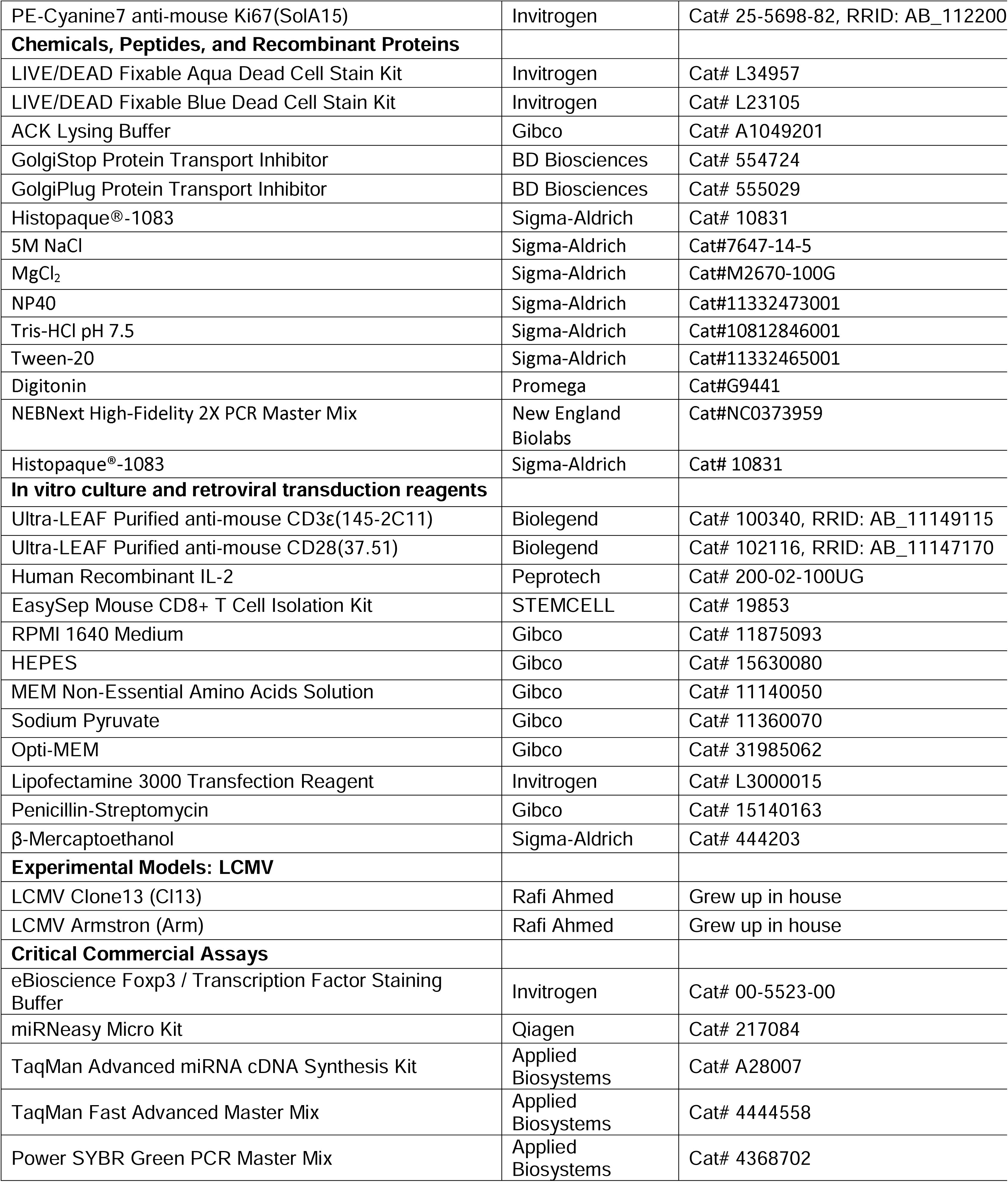
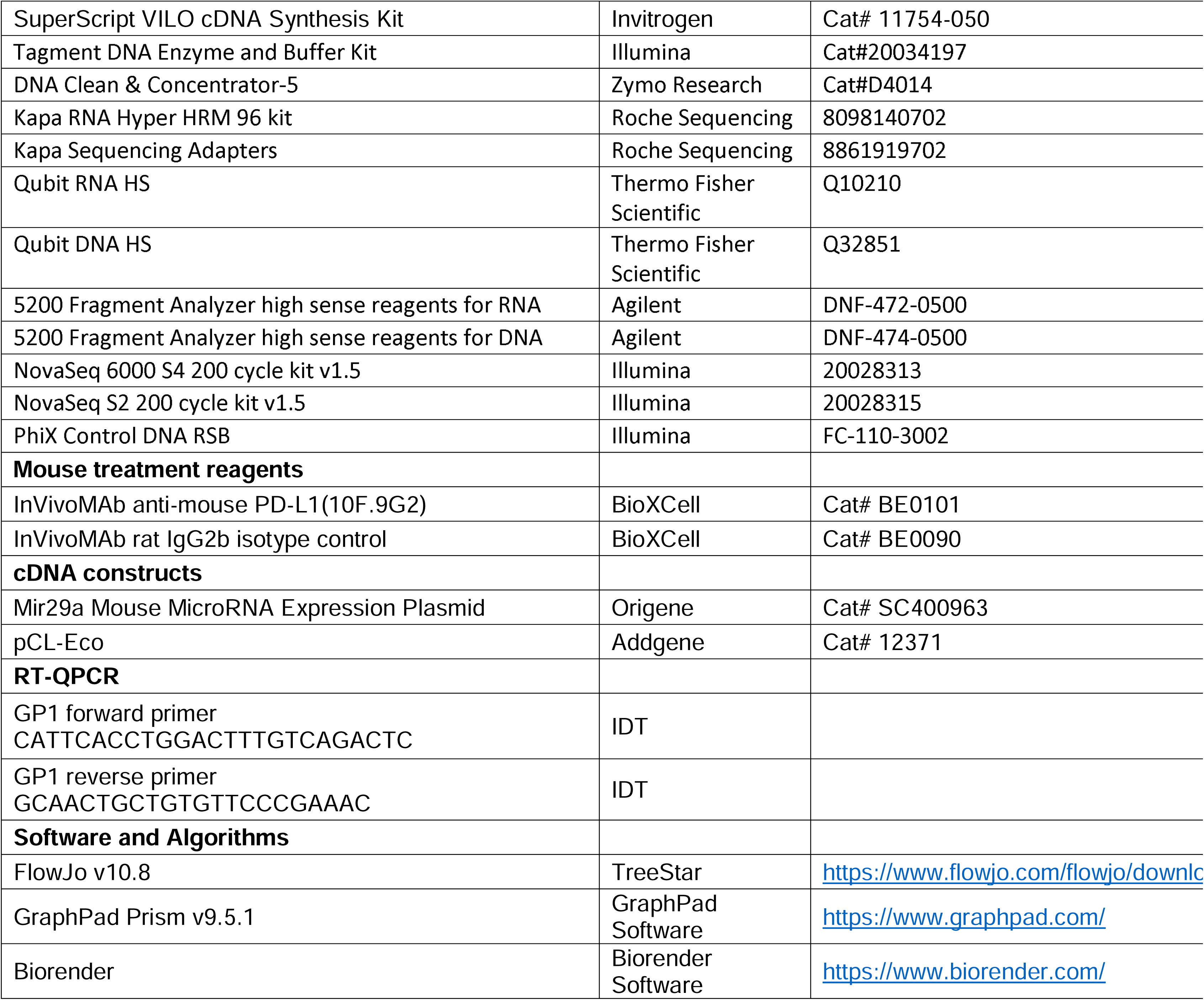

### Mice

C57BL/6J (CD45.2^+^ wild type, strain#: 000664) and B6.SJLPtprcaPepcb/BoyJ (CD45.1^+^ wild type, strain#: 002014) were purchased from Jackson Laboratories. P14 TCR transgenic mice expressing specific TCR for LCMV D^b^gp33-41 were kind gift from Dr. E. John Wherry (University of Pennsylvania) and bred in our facility. Both female and male 6 to 8-week-old mice were used as host for all the experiments in this study. In each experiment, the age and sex of mice were matched. All animal procedures were conducted in accordance with the Institutional Animal Care and Use Committee (IACUC) at the University of Miami.

### Virus and infection

Recipient mice were infected with 4 x 10^6^ LCMV clone 13 intravenously (i.v.). Quantitative real-time PCR was performed to test LCMV glycoprotein (GP1)-specific mRNA, using the forward primer 5’CATTCACCTGGACTTTGTCAGACTC3’ and the reverse primer 5’GCAACTGCTGTGTTCCCGAAAC3’. Rat IgG2b isotype (BioXcell) or aPD-L1 antibody (BioXcell) were administered intraperitoneally (200 µg/injection) every 3 days from day 21-33 p.i., a total of 5 injections.

### Cell preparations, *in vitro* T cell stimulation and transduction

Spleens were dissociated on 70-μM cell strainers and red blood cells were lysed using Ammonium-Chloride-Potassium (ACK) lysing buffer (Gibco). CD8 T cells were purified using magnetic separation (negative enrichment; STEMCELL) and stimulated with 100 U/mL recombinant human IL-2 in 48-well plates, which were precoated with 2.5 μg/mL anti-mouse CD3_ε_ and 10 μg/mL anti-mouse CD28 for 2 hrs. After 24-30 hrs of stimulation, T cells were transduced with miR-29a OE or control retroviral vectors in the presence of 8 μg/mL polybrene and 100 U/mL recombinant human IL-2 by spin infection at 2000g, 30°C for 1h (Kurachi et al., 2017; Stelekati et al., 2018). After 16 hrs, transduced cells were sorted for VEX^+^ or GFP^+^ and adoptively transferred to recipient mice. CD8 T cells were cultured *in vitro* using complete media: RPMI-1640 (Corning), 10% fetal bovine serum (ThermoFisher), 1% penicillin/streptomycin (Lonza), 0.1% β-mercaptoethanol (Sigma), 20mM HEPES (Gibco), 1mM Sodium Pyruvate (Gibco) and 100µM MEM Non-Essential Amino Acids (Gibco). To produce retroviruses, 293T cells (ATCC) were cultured with MSCV and pCL-Eco plasmids using Lipofectamine 3000 Transfection Reagent (Invitrogen). MiR-29a (MI0000576) overexpressing plasmid was obtained from Origene. miR-29a cDNA was cloned into MSCV-IRES-VEX and MSCV-IRES-GFP plasmid, as previously described (Stelekati et al., 2022).

### Flow cytometry and sorting

The following fluorochrome-conjugated antibodies and dilutions were used for flow cytometry and cell sorting. Antibodies from BD Biosciences: anti-CD39-BUV395 (Y23-1185, 1:100), anti-CD45.1-BUV615 (A20, 1:200), anti-CD45.2-BUV496 (104, 1:200), anti-CD69-BUV737 (H1.2F3, 1:500), anti-IL-2-APC (JES6-5H4, 1:100), anti-TCF-7/TCF-1-PE (S33-966, 1:100), BD Horizon™ Fixable Viability Stain 440UV (1:100). Antibodies from Biolegend: anti-CD244.2-APC (m2b4 (B6)458.1, 1:200), anti- CD127-PE-Cy5 (A7R34, 1:100), anti-CD160-PE (7H1, 1:100), anti-CD44-BV785 (IM7, 1:300), anti-CD45.1-AF700 (A20, 1:100), anti-CD45.2-APC-Cy7 (104, 1:200), anti-CD69-PE-Cy5 (H1.2F3, 1:100), anti-CD8a-BV650 (53-6.7, 1:400), anti-CX3CR1-APC/Fire 750 (SA011F11, 1:100), anti-CX3CR1-BV711 (SA011F11, 1:500), anti-IFN-γ-PE-Cy7 (XMG1.2, 1:100), anti-Ki-67-BV605 (16A8, 1:100), anti-KLRG1-BV605 (2F1/KLRG1, 1:200), anti-KLRG1-FITC (2F1/KLRG1, 1:200), anti-KLRG1-APC (2F1/KLRG1, 1:200), anti-CD223 (LAG3) -BV650 (11C3C65, 1:200), anti-Ly108-PB (330-AJ, 1:100), anti-PD-1-PE-Cy7 (29F.1A12, 1:200), anti-TNF-α-BV605 (MP6-XT22, 1:100). Antibodies from Invitrogen: LIVE/DEAD™ Fixable Aqua Dead Cell Stain Kit (1:100). Antibodies from Miltenyi Biotec: anti-TOX-APC (REA473, 1:100). Antibodies from Thermo Fisher Scientific: anti-CD39-PerCP-eF710 (24DMS1, 1:100), anti-EOMES-eFluor450 (Dan11mag, 1:200), anti-Granzyme B-PE-Texas Red (GB11, 1:50), anti-Ki67-PE-Cy7 (SolA15, 1:100).

Samples were stained with surface antibodies for 30 minutes at 4°C, washed and fixed. For intracellular staining, cell suspensions were fixed and permeabilized at 4°C for 30 minutes using Foxp3 Transcription Factor Staining Buffer Set (Thermo Fisher Scientific), followed by intracellular staining at 4°C for 60 minutes. To stain for intracellular cytokines, cells were cultured in 96-well plate for 5 hours at 37°C, 5% CO_2_ in complete media containing 0.4 µg/mL gp_33-41_ (NIH), GolgiPlug (BD Biosciences), GolgiStop (BD Biosciences), and anti-CD107a (Biolegend). Flow cytometry data were acquired on LSR-Fortessa-HTS and Cytek Aurora. Cells were sorted on BD FACS Aria Fusion. FACS data was analyzed using FlowJo software 10.8.

### RNA Isolation and Sequencing

RNA was isolated using Qiagen RNeasy Micro kit following manufacture’s protocol. For RNA sequencing, quality-control analysis, library generation, and RNA-seq were carried out by the Oncogenomics Core Facility at the University of Miami. RNA-seq libraries were prepared using the Roche Kapa RNA Hyper-Prep with Riboerase. RNA-sample quality was measured by Qubit. RNA integrity measured by Fragment Analyzer 5200. The library pool was sequenced on an Illumina NovaSeq 6000.

### DNA Isolation and ATAC library Preparation

5x10^4^ viable cells were resuspended in 1mL with cold RSB buffer and 0.01%D Digitonin, 0.1% NP40, and 0.1% Tween-20 for 3 minutes on ice, according to Omni-ATAC protocol (Corces et al., 2017). Cells were washed with RSB (without NP40 and digitonin) and nuclei were pelleted at 500 rcf for 10 min at 4°C. Cells were resuspended in transposition mixture and incubated at 37°C for 30 min in thermomixer with 1000 rpm. Transposase reagents were from Illumina. Cleanup reaction with Zymo DNA clean and concentrator-5 kit. Primer fragment ligation with PCR amplification was performed using NEBNext 2x Mastermix. Primers were purchased from IDT, primer sequences as previously reported (Buenrostro et al., 2015). Additional cycles required were determined with qPCR SYBR green (in DMSO). Library sequencing was carried out by the Oncogenomics Core Facility at the University of Miami. Libraries were sequenced on Illumina NovaSeq 6000.

### ATAC-seq primer sequences for library preparation

Ad1_noMX: AATGATACGGCGACCACCGAGATCTACACTCGTCGGCAGCGTCAGATGTG

Ad2.1_TAAGGCGA

CAAGCAGAAGACGGCATACGAGATTCGCCTTAGTCTCGTGGGCTCGGAGATGT

Ad2.2_CGTACTAG

CAAGCAGAAGACGGCATACGAGATCTAGTACGGTCTCGTGGGCTCGGAGATGT

Ad2.3_AGGCAGAA

CAAGCAGAAGACGGCATACGAGATTTCTGCCTGTCTCGTGGGCTCGGAGATGT

Ad2.4_TCCTGAGC

CAAGCAGAAGACGGCATACGAGATGCTCAGGAGTCTCGTGGGCTCGGAGATGT

Ad2.5_GGACTCCT

CAAGCAGAAGACGGCATACGAGATAGGAGTCCGTCTCGTGGGCTCGGAGATGT

Ad2.6_TAGGCATG

CAAGCAGAAGACGGCATACGAGATCATGCCTAGTCTCGTGGGCTCGGAGATGT

Ad2.7_CTCTCTAC

CAAGCAGAAGACGGCATACGAGATGTAGAGAGGTCTCGTGGGCTCGGAGATGT

Ad2.8_CAGAGAGG

CAAGCAGAAGACGGCATACGAGATCCTCTCTGGTCTCGTGGGCTCGGAGATGT

Ad2.9_GCTACGCT

CAAGCAGAAGACGGCATACGAGATAGCGTAGCGTCTCGTGGGCTCGGAGATGT

Ad2.10_CGAGGCTG

CAAGCAGAAGACGGCATACGAGATCAGCCTCGGTCTCGTGGGCTCGGAGATGT

Ad2.11_AAGAGGCA

CAAGCAGAAGACGGCATACGAGATTGCCTCTTGTCTCGTGGGCTCGGAGATGT

Ad2.12_GTAGAGGA

CAAGCAGAAGACGGCATACGAGATTCCTCTACGTCTCGTGGGCTCGGAGATGT

Ad2.13_GTCGTGAT

CAAGCAGAAGACGGCATACGAGATATCACGACGTCTCGTGGGCTCGGAGATGT

Ad2.14_ACCACTGT

CAAGCAGAAGACGGCATACGAGATACAGTGGTGTCTCGTGGGCTCGGAGATGT

Ad2.15_TGGATCTG

CAAGCAGAAGACGGCATACGAGATCAGATCCAGTCTCGTGGGCTCGGAGATGT

### Bioinformatic Analysis

Raw FASTQ from RNAseq paired-end sequencing were processed with cutadapt (v5.1) using parameters –adapter AGATCGGAAGAG -A AGATCGGAAGAG --minimum-length 10. Reads were mapped to the GRCm39 mouse genome with GENCODE vM35 basic annotations using STAR (v2.7.11.b) and RSEM (v1.3.1). Counts were extracted and differential expression analysis was performed with DESeq2 (v1.49.3). A threshold of absolute log_2_ fold change and 5% FDR was used to identify significant genes. Log_2_-normalized FPKM values were used for Significance Analysis of Microarrays (SAM) analysis (samr v3.0). The observed SAM score was used as input for GSEA (v4.4.0) with annotations from KEGG, Hallmark, Reactome gene sets; significant pathways were defined as those with FDR<10%. Gene signatures from publicly available datasets were used as reference datasets: T_EX_ (Bengsch et al., 2018), T_EX_ vs T_MEM_ (GSE9650), T_EX_(PD-1^+^CD39^+^)(Giles et al., 2022), Terminal T_EX_ (Yao et al., 2019), Progenitor T_EX_ (Yao et al., 2019), T_EFF_ (Milner et al., 2020), T_EFF_ vs T_MEM_(GSE1000002_1582_200_UP), T_EFF_ vs T_EX_ (GSE9650), T_MEM_ (Milner et al., 2020), T_MEM_ vs T_EFF_ (GSE10239), T_CM_ vs T_EM1_ (Giles et al., 2022), T_CM_ vs T_EM2_ (Beltra et al., 2020), Progenitor T_MEM_ (Yao et al., 2019), T_RM_(Milner) (Crowl et al., 2022), T_RM_(Mackay) (Crowl et al., 2022), T_RM_(Crowl) (Crowl et al., 2022), circulating T cells (Milner et al., 2017), miR-29a targets from TargetScan (McGeary et al., 2019) (v7.2) as well as our previous miR-29a signatures (Stelekati et al., 2022). Protein-protein interactions were drawn from InAct(Del Toro et al., 2022) (Accessed Sept 20^th^, 2025) and visualized using cytoscape (Shannon et al., 2003).

ATAC-seq data analysis was performed by running nf-core/atacseq pipeline(v1.1.2), which includes read trimming with Trim Galore (v0.6.7) using Cutadapt (v3.4) (Phred quality cutoff 20; Nextera transposase adapter sequence ‘CTGTCTCTTATA’), alignment to the GRCm39 genome using bwa (v0.7.17-r1188), duplicate removal with Picard (v3.0.0) and peak calling with MACS2 (v2.2.7.1). Peak annotation was manually performed at overlapping genes using gencode(Mudge et al., 2025) vM35. Differential peaks were called with DESeq2 with a threshold of 5% FDR and an absolute log_2_ fold change > 1. Motif enrichment analysis was performed with HOMER (v5.1). Differential peaks were intersected with genomic coordinates of genes (extended by ±1,000bp) and enrichment of T cell expression signatures as well as Reactome pathways were assessed using hypergeometric tests, with P-values adjusted to FDR. Differential peaks were also intersected from previous studies (Abdel-Hakeem et al., 2021; Pauken et al., 2016), and significant overlaps were evaluated using hypergeometric tests with FDR correction.

### Statistical Analysis

Flow cytometry data were analyzed using GraphPad Prism V.9 software. Statistical analysis was presented as mean ± SEM. Mann-Whitney test was used in two-group analyses. One-way analysis of variance (ANOVA) was used in three or more group analyses, with Tukey’s multiple comparisons test. Statistical significance is indicated by * = P<0.05, ** = P<0.01, and *** = P<0.001.

**Figure S1.**
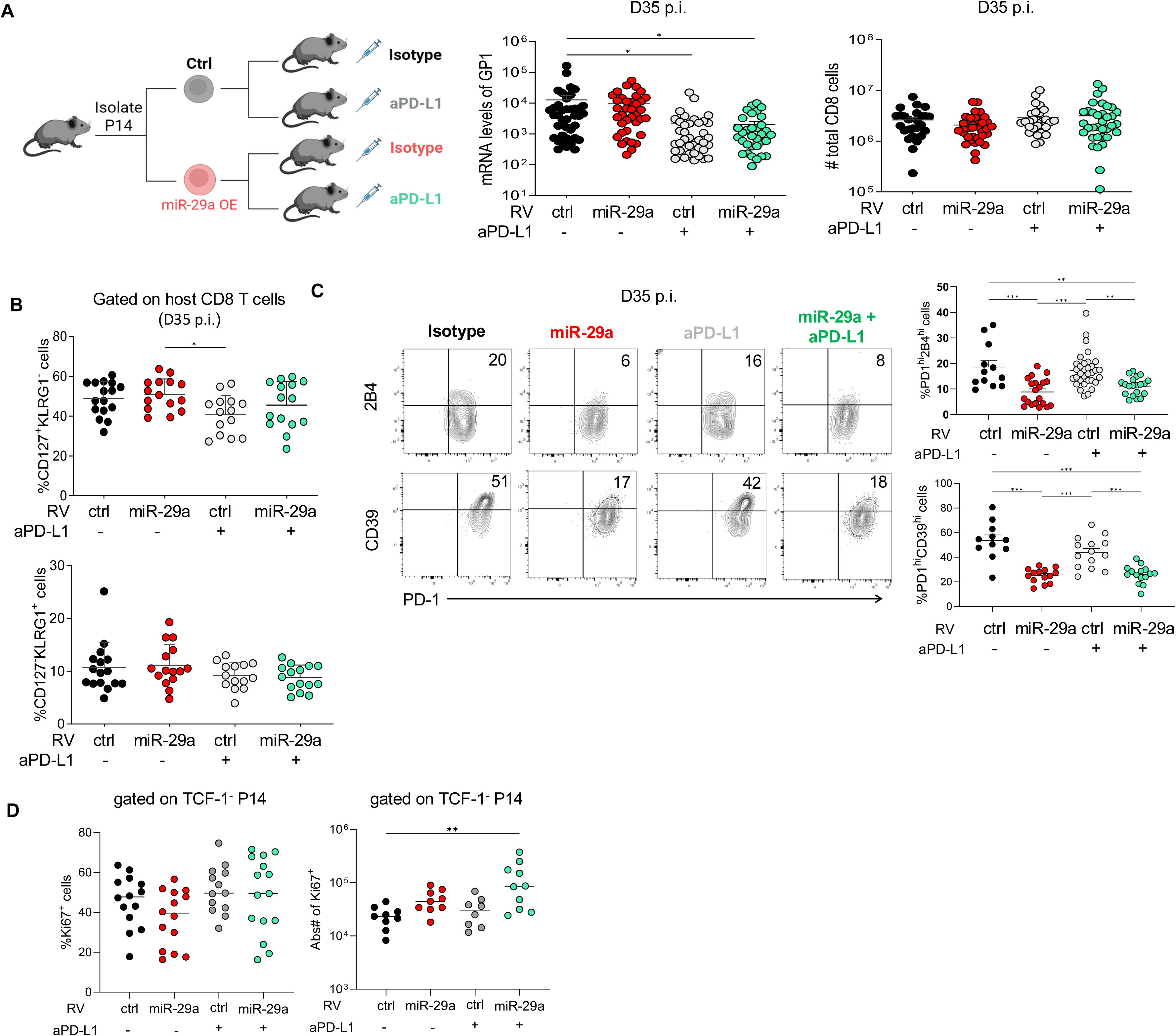
MiR-29a OE does not regulate proliferation in terminal T_EX_. **(A)** P14 cells were transduced with miR-29a OE or control retrovirus, sorted for VEX^+^ expression and transferred into congenically marked recipient mice that were infected with LCMV cl-13 at 48 hrs earlier. Isotype or anti-PD-L1 (200 µg/injection) was administered every third day starting at d21 p.i., for a total of 5 doses. LCMV GP1 gene was quantified by qPCR at day 35 p.i. Absolute numbers of CD8 T cells at day 35 p.i. **(B)** Frequencies of CD127^+^KLRG1^-^and CD127^-^KLRG1^+^ T cells in host CD8 T cells from Figure 1A**. (C)** Frequency of PD-1^hi^2B4^hi^ and PD-1^hi^CD39^hi^ cells in donor P14 cells from Figure 1A. FACS plots are gated on donor VEX^+^ P14 cells and numbers represent percentages out of total donor VEX^+^ P14 cells. **(D)** Ki- 67 expression was evaluated in TCF-1^-^ P14 cells. Data are pooled from 5 independent experiments with n>9. Results are indicated as mean ± SEM. Statistical analysis was performed using One-way ANOVA with Tukey’s multiple comparisons. (\**P* < 0.05, \*\**P* < 0.01, and \*\*\**P* < 0.001). Ctrl, control RV.

**Figure S2.**
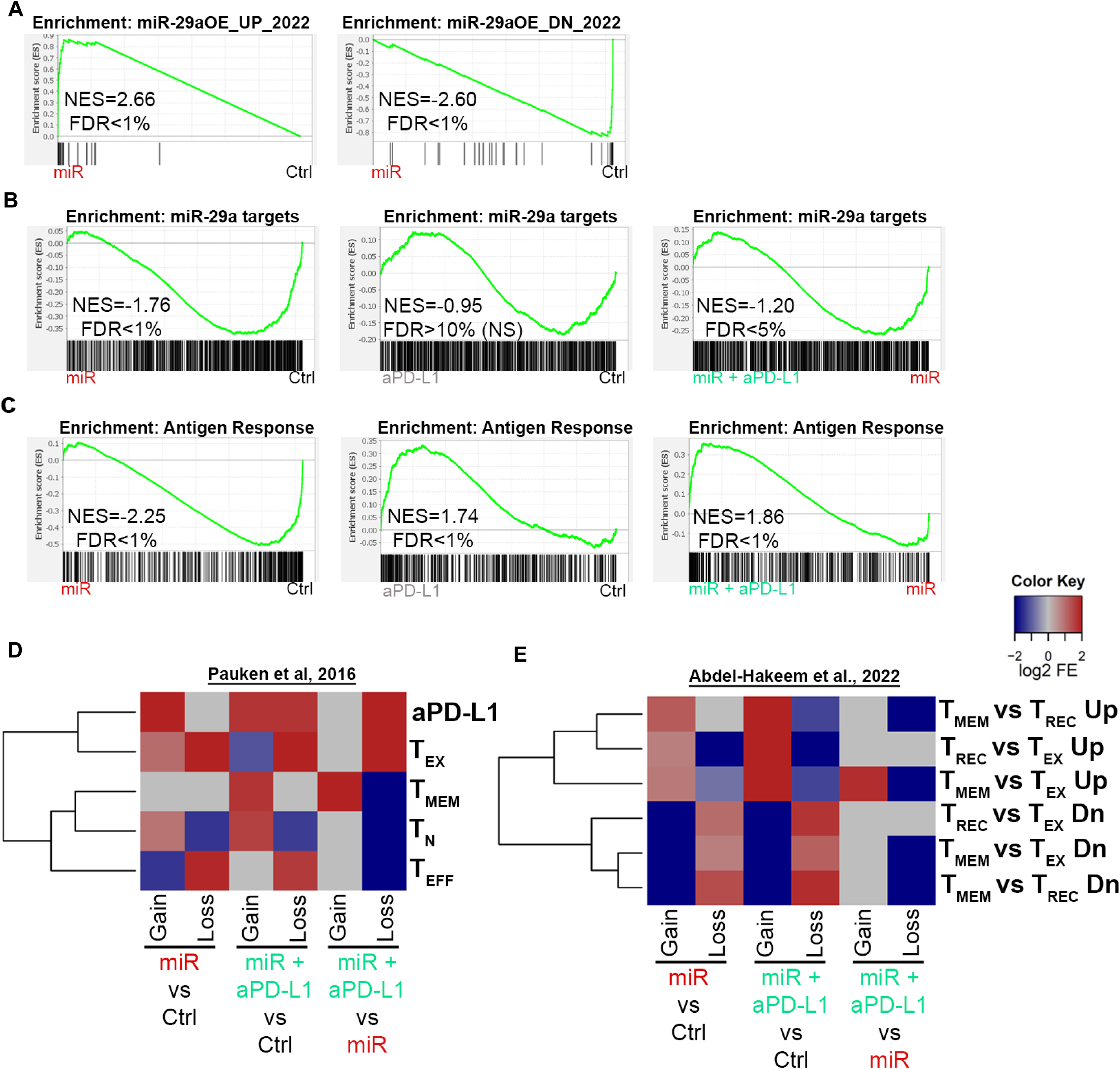
MiR-29a OE downregulates transcriptional and epigenetic signatures of chronic antigen stimulation. P14 cells were transduced with miR-29a OE or control retrovirus, sorted for VEX^+^ expression and transferred into congenically marked recipient mice that were infected with LCMV cl-13 at 48 hrs earlier. Isotype or aPD-L1 (200 µg/injection) was administered every third day starting at d21 p.i., for a total of 5 doses. VEX^+^ P14 cells were sorted from spleens at d35 p.i. and RNA Sequencing and ATAC Sequencing were performed. **(A)** GSEA of miR-29a OE vs control signatures using our previously published dataset (Stelekati et al., 2022). **(B)** GSEA using miR-29a predicted mRNA targets from TargetScan. **(C)** GSEA using dataset on Antigen Response (Goldrath et al., 2004). **(D-E)** Heatmaps of overlapped peak region accessibility against differentiated T cell types (Abdel-Hakeem et al., 2021; Pauken et al., 2016).

**Figure S3.**
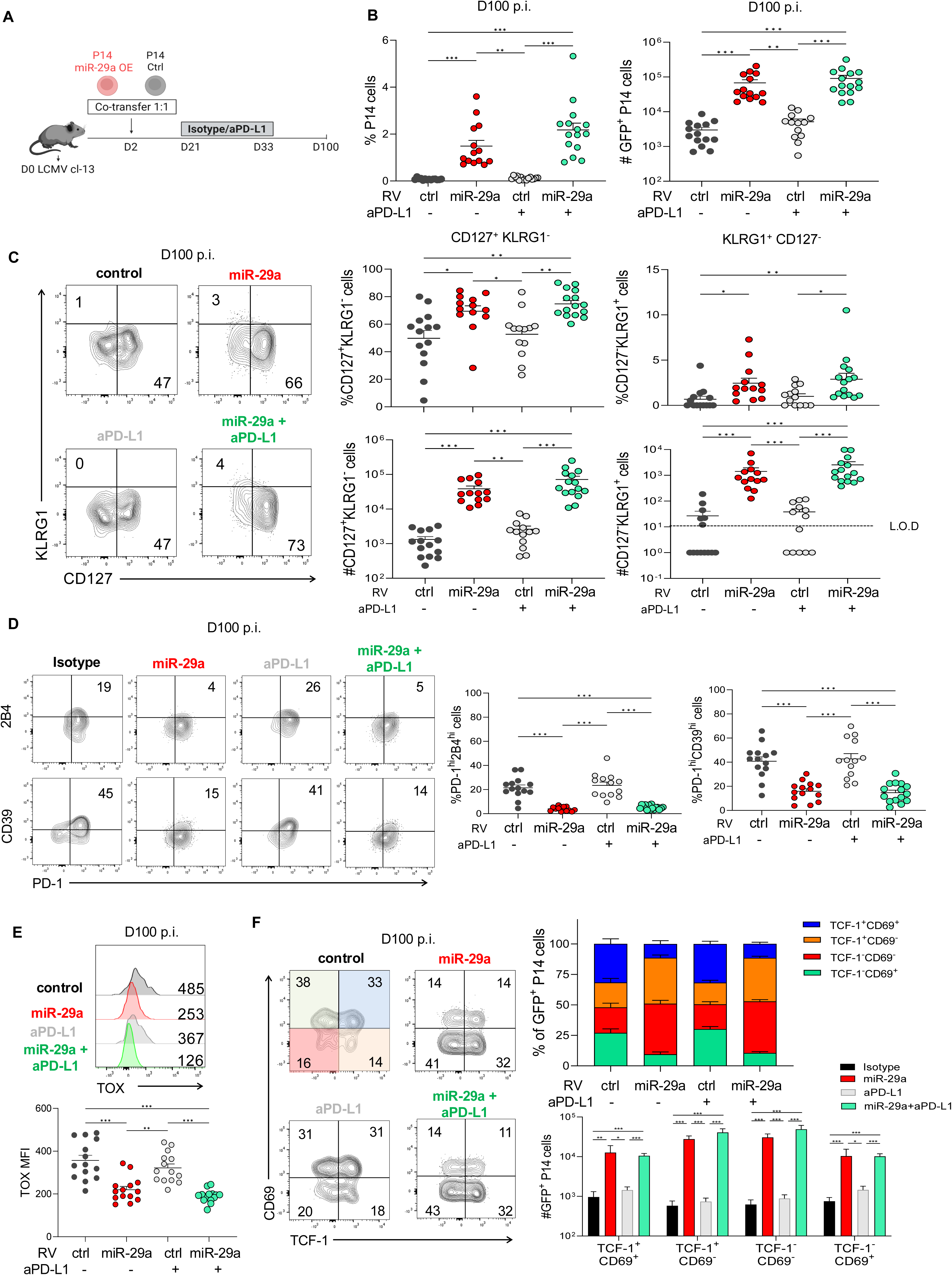
MiR-29a OE attenuates T cell exhaustion long-term. **(A)** P14 cells were transduced with miR-29a OE or control retrovirus, sorted for GFP^+^ expression and co-transferred into host mice at a 1:1 ratio. The host mice had been infected with LCMV cl-13 2 days prior to transfer. Isotype or aPD-L1 (200 µg/injection) were administered as described above. **(B)** Quantification of donor P14 cells in the spleens at d100 p.i. **(C)** Frequencies of CD127^+^KLRG1^-^ and CD127^-^KLRG1^+^ P14 cells at d100 p.i. **(D)** Frequencies of PD-1^hi^2B4^hi^ and PD-1^hi^CD39^hi^ P14 cells at d100 p.i. **(E)** Histogram of TOX expression P14 cells. **(F)** Frequencies of 4 T_EX_ subsets. Data are pooled from 2 independent experiments with n > 5. FACS plots are gated on donor GFP^+^ P14 cells and numbers represent percentages out of total donor GFP^+^ P14 cells. Results are indicated as mean ± SEM. Statistical analysis was performed using Kruskal-Wallis test with Dunn’s multiple comparisons (\**P* < 0.05, \*\**P* < 0.01, and \*\*\**P* < 0.001). Ctrl, control RV. Illustration in **(A)** created using BioRender.com.

**Figure S4.**
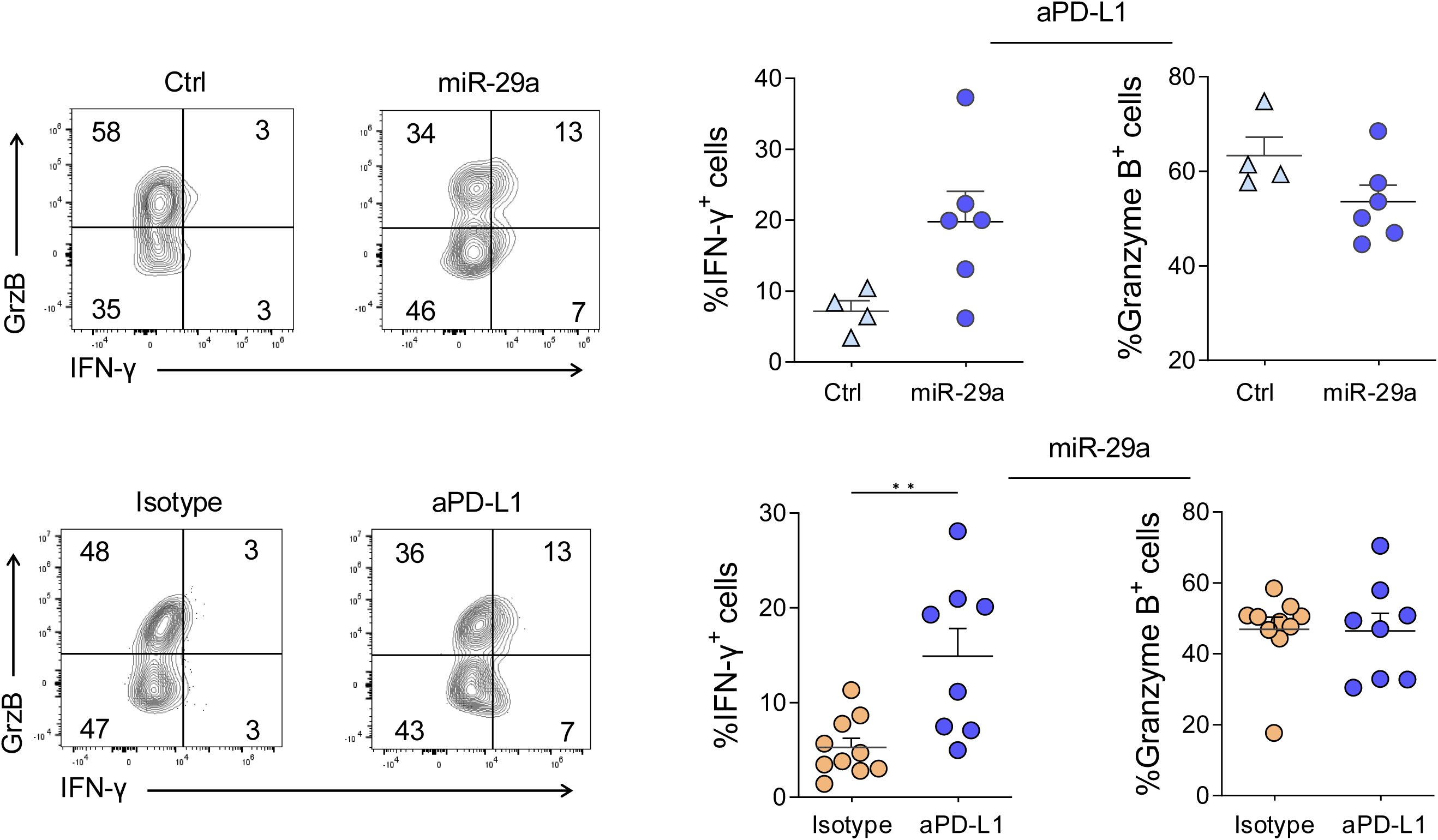
Addition of aPD-L1 increases cytolytic molecule production in miR-29a OE progenitor T_EX_. Frequencies (top) of IFN-γ^+^ and Granzyme B^+^ P14 cells from Figure 4C. Frequencies (bottom) of IFN-γ^+^ and Granzyme B^+^ P14 cells from Figure 4G. FACS plots are gated on donor GFP^+^ P14 cells and numbers represent percentages out of total donor GFP^+^ P14 cells.

**Table S1. MiR-29a OE regulates the transcriptional and epigenetic signatures of CD8 T cells during chronic antigen stimulation.**

P14 cells were transduced with miR-29a OE or control retrovirus, sorted for VEX^+^ expression and transferred into congenically marked recipient mice that were infected with LCMV cl-13 at 48 hrs earlier. Isotype or aPD-L1 (200 µg/injection) was administered every third day starting at d21 p.i., for a total of 5 doses. VEX^+^ P14 cells were sorted from spleens at d35 p.i. and RNA Sequencing and ATAC Sequencing were performed. Table contains all differentially expressed genes (DEG) and differentially accessible regions (DAR) among pairwise comparisons, and pathways differentially enriched in pairwise comparisons.

